# Development of Dry Powder Formulations of AS01_B_ containing vaccines using Thin-Film Freeze-Drying

**DOI:** 10.1101/2022.02.27.482135

**Authors:** Khaled AboulFotouh, Haiyue Xu, Robert O. Williams, Zhengrong Cui

## Abstract

AS01_B_ is a liposomal formulation of two immunostimulants namely 3-O-desacyl-4’-monophosphoryl lipid A (MPL) and QS-21. The liposomal formulation of AS01_B_ reduces the endotoxicity of MPL and the lytic activity of QS-21; however, it renders the adjuvant sensitive to accidental slow freezing. The liposomal formulation also represents a major challenge towards the formulation of dry powders of vaccines containing AS01_B_. In the present study, we tested the feasibility of applying thin-film freeze-drying (TFFD) to engineer dry powders of the AS01_B_ liposomal adjuvant alone or vaccines containing AS01_B_ as an adjuvant. Initially, we showed that after the AS01_B_ liposomal adjuvant was subjected to TFFD using sucrose as a stabilizer at 4% *w/v*, the particle size distribution of AS01_B_ liposomes reconstituted from the dry powder was identical to the liquid adjuvant before drying. We then showed using ovalbumin (OVA) as a model antigen adjuvanted with AS01_B_ (AS01_B_/OVA) that subjecting the AS01_B_/OVA vaccine to TFFD and subsequent reconstitution did not negatively affect the AS01_B_ liposome integrity, nor the immunogenicity of the vaccine. Importantly, the thin-film freeze-dried vaccine was not sensitive to repeated freezing-and-thawing. Finally, the feasibility of using TFFD to prepare dry powders of AS01_B_-adjuvanted vaccines was further confirmed using AS01_B_-adjuvanted Fluzone Quadrivalent and Shingrix, which contains AS01_B_. It is concluded that the TFFD technology can enable the formulation of AS01_B_-adjuvanted vaccines as freezing-insensitive dry powders in single-vial presentation.

## 1. Introduction

Many of the human vaccines approved by the Food and Drug Administration (FDA) for use in the United States contain an adjuvant or an adjuvant system [1]. Adjuvants are included in vaccines to enhance their immunogenicity [2]. Vaccine adjuvants include immunopotentiators that directly activate antigen-presenting cells (APCs) *via* the pattern-recognition receptors (*e*.*g*., Toll-Like Receptors (TLRs)) and delivery systems that help localize and target the antigenic components to the APCs [3]. Aluminum salts (*i*.*e*., aluminum (oxy)hydroxide) are the most commonly utilized vaccine adjuvants. They are present in ~ 77% of FDA-approved adjuvanted vaccines and were the only adjuvants employed in licensed human vaccines for close to a century [1, 4, 5]. However, in the past 20 years five new adjuvants were introduced in FDA-approved human vaccines, namely AS03, AS04, MF59^®^, CpG 1018, and AS01_B_ [6].

GlaxoSmithKline has developed a series of adjuvant systems including adjuvant system (AS) 01 and AS02 to help antigens elicit durable humoral and cellular immune responses [7]. Among these systems, AS01_B_ induced the strongest and the most persistent humoral and CD4^+^ T-cell responses against recombinant protein antigens *via* stimulating massive production of efficient APCs (*e*.*g*., dendritic cells) [7-9]. AS01_B_ is a liposomal formulation of two immunostimulants that work synergistically namely 3-O-desacyl-4’-monophosphoryl lipid A (MPL) and QS-21 [5]. MPL is the detoxified derivative of lipopolysaccharide from the Re595 strain of *Salmonella minnesota* and activates the innate immune responses through direct activation of TLR4-expressing APCs [5, 7]. QS-21 is a highly purified triterpene glucoside (*i*.*e*., fraction 21) extracted from the bark of *Quillaja saponaria* Molina [5, 10]. In mice, QS-21 activates the nucleotide binding and oligomerization domain (NOD)-like receptor P3 inflammasome complex present within the cytosol of APCs and thus optimizes the T-cell response [11]. The durable humoral and cellular immune responses induced by AS01_B_-adjuvanted vaccines led to the recommendation of its utilization as an adjuvant in vaccine candidates for protection against complex pathogens (*e*.*g*., *Plasmodium falciparum*) [7]. In 2017, Shingrix^®^ vaccine (*i*.*e*., Zoster Vaccine Recombinant, Adjuvanted, suspension for intramuscular injection) was approved by the FDA as the first AS01_B_-adjuvanted vaccine for human use.

The liposomal formulation of AS01_B_ reduces the endotoxicity of MPL and the lytic activity of QS-21 while maintaining their adjuvant activity; however, the liposomal formulation also renders the adjuvant sensitive to slow accidental freezing [7, 12]. Liposomes have been employed as nanocarriers in several FDA-approved pharmaceutical products; however, the promise of liposomes as drug delivery vehicles is challenged by their physical (*e*.*g*., drug leakage and aggregation or fusion) and chemical (*e*.*g*., lipid oxidation and hydrolysis) instability [13-15]. Thus, the AS01_B_ adjuvant of Shingrix vaccine is provided as a suspension in a separate vial, which is used to reconstitute the lyophilized varicella zoster virus glycoprotein E (VZV-gE) antigen immediately before administration. The AS01_B_ suspension and lyophilized VZV-gE antigen should be stored refrigerated between 2°C and 8°C and should not be frozen. The vaccine should be discarded if the adjuvant suspension and/or the lyophilized antigen have been accidentally frozen or exposed to temperatures > 8ºC. There are reports of vaccines being exposed to freezing temperatures during shipping or storage, which could result in a significant loss of freezing-sensitive vaccines (*e*.*g*., AS01_B_-adjuvanted vaccines) [16]. As such, improved formulations of AS01_B_-adjuvanted vaccines that permit less stringent storage conditions and greater overall stability would be highly advantageous.

Although drying of vaccines is overall challenging, it is the most efficient approach to enhance the stability of liposomal products [14]. Freeze-drying has been extensively explored to formulate liposomal dry powders, particularly for inhalation [17-20]. Unfortunately, freeze-drying was reported to significantly affect the liposome structure (*e*.*g*., transition from unilamellar to bi- or multilamellar vesicles or bilayer swelling) and particle size distribution after reconstitution [14, 21, 22]. For instance, the damage of lipid bilayer integrity during freeze-drying resulted in severe leakage of encapsulated ciprofloxacin [13]. These events can be attributed to the compressive stress of the bilayer resulting from the reduction of spacing between the phospholipid head groups during freezing [15]. Thus, various formulation approaches have been explored to protect the liposomes against freeze-drying stress. These approaches include the addition of a support (*e*.*g*., gelatin) in the liposome interior compartment [14], or coating of the liposomes with a polymer (*i*.*e*., calcium alginate) to enhance their physical stability [23]. Additionally, conventional shelf freeze dried liposomes cannot be readily inhaled [24]. The powder requires further processing such as the addition of carriers (*e*.*g*., SorboLac 400 and sieved Pharmatose 325 M) [18, 19], sieving [20], or micronization [25] in order to enhance its aerosol performance. The extra processing step not only extends the formulation time but also affects the activity of the active ingredients in the liposomes. For instance, β-glucuronidase content and activity in a liposomal formulation were reduced to 43% and 29% after lyophilization, respectively [25]. Micronization further reduced the enzyme content and activity to 36% and 22%, respectively, compared to the freshly prepared liposomes [25]. Ultimately, the need for additional excipients may represent a challenge for regulatory approval. Other technologies such as spray-drying [26, 27] and spray freeze-drying [13, 28, 29] have also been explored to formulate liposomal dry powders. However, the temperature of drying gas and/or the high shearing forces in spray-drying and spray freeze-drying processes can result in disruption of the liposomal bilayer structure [22, 30]. Consequently, the mean particle size of dry liposomes prepared using these drying processes increased following reconstitution [13, 26]. The liposome particle size is a key attribute that affects the body’s immune response to the loaded antigens [31]. Liposomes having a mean particle size comparable to that of virus particles can significantly enhance the antigen uptake by APCs [32]. The liposome particle size also determines the residence time of liposome in the cells of the lymph nodes and thus in the activation of innate immune responses [33]. Other physical and structural properties of liposomes such as lamellarity, surface charge, and bilayer fluidity also affect the elicited immune responses [34]. Thus, the liposome physical properties should be maintained after drying and reconstitution. Thus, complex formulation approaches (*e*.*g*., addition of internal and external lyoprotectants or a high concentration of stabilizing agent mixture) may be required to enhance the stability of liposomes against the stresses [13, 26].

Ultimately, an ideal drying technology should maintain the physical structure and the physicochemical properties of liposomes (*e*.*g*., mean particle size and zeta potential) [14]. In this study, we hypothesized that thin-film freeze-drying (TFFD) technology can be used to formulate dry powders of the AS01_B_ liposomes, alone or mixed with split virus or subunit protein antigens, while maintaining the liposome integrity and immunogenicity of the antigenic proteins upon reconstitution. In thin-film freezing (TFF), the liquid formulations of small molecules (*e*.*g*., remdesivir) [35] or large molecules (*e*.*g*., proteins, siRNA, virus-like particles or monoclonal antibodies) [36-39] are frozen into thin films rapidly, followed be sublimation to remove the solvent [40, 41]. The TFF process achieves cooling rates (*i*.*e*., 100-100 K/s) intermediate between spray freeze-drying and conventional shelf freeze-drying, which are sufficient to prevent particle growth during freezing [42]. The ultrafast cooling in the TFF process results in the formation of small ice crystals and a homogeneous distribution of the stabilizing excipients, which might reduce the disruption of the liposomal bilayer structure [30]. However, slow freezing is also advantageous in terms of reducing the osmotic pressure, minimizing the formation of ice crystals in the inner aqueous compartment, and providing more time for liposomes to recover from deformations caused by osmotic pressure and mechanical stress [15]. The gas-liquid interface of TFF process is much smaller than the spray freeze-drying process which in turn results in minimal protein denaturation [42]. Interestingly, TFF produces low density, brittle matrices with desired aerosol performance properties for respiratory delivery 91 [43].

Here, TFFD was applied to convert liquid formulations of AS01_B_ or AS01_B_-adjuvanted vaccines into dry powders with a low concentration of a single stabilizing excipient, while maintaining the particle size distribution, adjuvanticity of the AS01_B_, as well as the immunogenicity of the vaccines. The developed adjuvant and vaccine dry powders are not sensitive to accidental slow freezing. Ultimately, the TFFD technology can contribute to the cost reduction of vaccination programs and increase vaccination rate by allowing single-vial presentation of vaccines containing the liposomal AS01_B_ as an adjuvant by reducing the production cost and the wastage of vaccine doses exposed to accidental freezing. The dry powder vaccines may also be stored free of the cold chain and administered *via a* noninvasive route (*e*.*g*., intranasal or pulmonary).

## 2. Materials and methods

### 2.1 Chemicals

1,2-dioleoyl-sn-glycero-3-phosphocholine (DOPC) and cholesterol were from Avanti Polar lipids (Alabaster, AL). Ovalbumin (OVA), 3-O-desacyl-4’-monophosphoryl lipid A (MPL), trehalose dihydrate and D-mannitol were from Sigma Aldrich (St. Louis, MO). QS-21 was from Desert King International (San Diego, CA). Sucrose was from Merck KGaA (Darmstadt, Germany).

### 2.2 In house preparation of AS01_B_ adjuvant and AS01_B_-adjuvanted OVA model vaccine

AS01_B_ was prepared in house as previously described with modifications [7]. Briefly, the liposome formulation was prepared by dissolving 1 mg DOPC, 0.25 mg of cholesterol, and 50 µg of MPL in 2 mL ethanol. A lipid film was formed by solvent evaporation under a gentle stream of nitrogen gas. The lipid film was then hydrated with phosphate buffered saline (PBS, 10 mM, pH 7.2) to form liposomes. QS-21 (50 µg) in aqueous solution was added to the liposomes and the volume was adjusted to 0.5 mL using PBS (10 mM, pH 7.2). AS01_B_-adjuvanted OVA (AS01_B_/OVA) model vaccine was prepared by mixing OVA, dissolved in PBS, with AS01_B_ at a concentration of 50 µg OVA/0.5 mL.

### 2.3 Preparation and characterization of dry powders of AS01_B_ and AS01_B_/OVA model vaccines

#### 2.3.1 Thin-film freeze-drying

Liquid AS01_B_ adjuvant and AS01_B_/OVA vaccines were converted to dry powders using TFFD. 124 Sucrose was employed as a stabilizer in liquid AS01_B_ formulation at the same concentration level as in the 125 lyophilized VZV-gE antigen of the Shingrix^®^ vaccine (*i*.*e*., total lipids to sucrose ratio of 1:15 *w/w*, 126 respectively). Liquid AS01_B_/OVA vaccine formulations were prepared using sucrose, trehalose dihydrate, 127 or D-mannitol as a stabilizer at total lipids to stabilizer ratio of 1:4, 1:8, 1:15 or 1:30 *w/w*, respectively. 128 First, the liquid adjuvant or vaccine formulations were frozen into thin films by single-vial TFF as 129 previously described [35, 44]. A cryogenically cooled surface was created by immersing a wide-mouth 130 glass vial (Fisher brand) in liquid nitrogen for ∼10 min. Then, liquid formulations (500 µL) were 131 individually dropped onto the inner wall of the vial bottom using a 21-gauge syringe so that the liquid 132 droplets were rapidly frozen into thin-films. Thin-films of AS01_B_/OVA vaccine with sucrose at a ratio of 133 1:15 *w/w* (lipids : sucrose) were also prepared by bulk TFF. Briefly, AS01_B_/OVA vaccine formulations 134 were dropped onto a rotating, cryogenically cooled stainless-steel drum to form frozen thin-films. The 135 temperature of drum surface was maintained at -50°C or -180°C. The frozen thin-films were collected in 136 liquid nitrogen and then maintained in -80°C freezer until lyophilization using a VirTis Advantage bench top tray lyophilizer (The VirTis Company, Inc. Gardiner, NY). Lyophilization was performed over 60 h at pressures ≤ 100 mTorr. The shelf temperature was maintained at -40°C for 20 h and then gradually ramped to +25°C, over 20 h. Throughout the secondary drying phase, the vials were kept at +25°C for additional 20 h. The vials were then stored at room temperature for future use.

#### 2.3.2 In vitro characterization of AS01_B_ and AS01_B_/OVA vaccine dry powder

Dry powders of AS01_B_ adjuvant or AS01_B_/OVA vaccine were reconstituted in water, and particle size distribution, polydispersity index (PDI) and zeta potential values were meausred using a Malvern Zeta Sizer Nano ZS (Worcestershire, UK) after dilution with milli-Q water. The integrity of OVA protein in the reconstituted AS01_B_/OVA vaccine and its liquid counterpart was investigated using SDS-PAGE analysis and intrinsic tryptophan fluorescence spectometry. For SDS-PAGE analysis, 10 µL of reconstituted AS01_B_/OVA vaccine was mixed with Laemmli Sample Buffer (Bio-Rad, Hercules, CA) and β-mercaptoethanol (2% *v/v*, Sigma-Aldrich). Samples were heated at 95°C for 5 min prior to loading onto 4-20% Mini-PROTEAN^®^ TGX™precast polyacrylamide gel (Bio-Rad). Gel electrophoresis was done at 100 V for 90 min. Gel was stained in a Bio-Safe™ Coomassie G-250 Stain (Bio-Rad). The intrinsic fluorescence of tryptophan residues in OVA protein was recorded using a PTI QuantaMaster Spectrofluorometer (Photon Technology International, Santa Clara, CA) at an excitation wavelength of 295 nm and the emission spectrum was collected from 300 nm to 400 nm [45]. Powder crystallinity was evaluated using a Rigaku Oxford Diffraction HyPix6000E Dual Source diffractometer (Tokyo, Japan) using a μ-focus sealed tube Cu Ka radiation source (λ = 1.5418Å) with collimating mirror monochromators. The instrument was operated at an accelerating voltage of 50 kV at 0.8 mA. The data were collected at 100 K using an Oxford Cryostream low temperature device (Oxford Cryosystems Ltd, Oxford, United Kingdom). The data collection and data reduction were performed using Rigaku Oxford Diffraction’s CrysAlisPro V 160 1.171.42.25a.

#### 2.3.3 Repeated freezing and thawing

To investigate the effect of repeated freezing and thawing on the mean particle size of thin-film freeze-dried powder of AS01_B_/OVA vaccine and its liquid counterpart, liquid formulation (0.5 mL) and dry powder were subjected to three consecutive cycles of freezing at -20°C for 8 h and thawing at 4°C for 16 h. At the end of the third cycle, the powder was reconstituted and adequately diluted with milli-Q water and the Z-average hydrodynamic particle size was determined by dynamic light scattering (DLS).

#### 2.3.4 Animal studies

The protocol of animal study was approved by the Institutional Animal Care and Use Committee at The University of Texas at Austin. Female C57BL/6 mice (6-8 weeks old) were from Charles River Laboratories (Wilmington, MA). The mice were allowed free access to food and water. Mice (five per group) were subcutaneously injected twice, three weeks a part with AS01_B_/OVA vaccine reconstituted from dry powder, liquid AS01_B_/OVA vaccine that was not subjected to TFFD, PBS (10 mM, pH 7.2) or OVA solution in PBS. The dose of OVA was 5 µg/mouse/dose while the dose of AS01_B_ was 135 µg/mouse/dose (*i*.*e*., 100 µg of DOPC, 25 µg of cholesterol, 5 µg of MPL and 5 µg of QS-21). Mice were euthanized two weeks after the second immunization to collect blood and spleens. Blood samples were centrifuged at 14,000 rpm for 10 min to collect sera. Antibody responses including specific IgG, IgG1 and IgG2a in mouse sera were measured using an ELISA assay as previously described [46]. Splenocyte proliferation assay was performed as previously described [47]. Briefly, spleens were isolated and homogenized. Splenocytes were seeded into 24-well tissue culture plates at 2×10^6^ cells/well in RPMI1640 medium supplemented with 0.05 mM ß-mercaptoethanol and 10% FBS. Splenocytes stimulated with OVA at a final concentration of 10 µg/mL of culture medium and unstimulated cells served as positive and negative controls of the proliferation assay, respectively. The plates were incubated at 37ºC in a humidified atmosphere with 5% CO_2_ for 48 h. Cell proliferation was evaluated using an MTT assay. Proliferation index was calculated as the proliferation of cells stimulated with OVA divided by the proliferation of unstimulated cells. The culture supernatants of stimulated cells were collected for cytokine (*i*.*e*., IL-4 and IFN-γ) assays using ELISA kits (BioLegend, CA).

### 2.4 Preparation and characterization of dry powders of AS01_B_-adjuvanted Fluzone Quadrivalent^®^ vaccine

AS01_B_-adjuvanted Fluzone Quadrivalent (AS01_B_/FluzoneQuad) was prepared by mixing 0.5 mL of inhouse prepared AS01_B_ adjuvant with an equivalent volume of Sanofi Pasteur’s Fluzone Quadrivalent (FluzoneQuad) (from UT Austin, Forty Acres Pharmacy). Sucrose was employed as a stabilizer at sugar to total lipid ratio of 15:1 *w/w*. Then, liquid formulation of AS01_B_/FluzoneQuad vaccine was frozen into thin-films by single-vial TFF followed by lyophilization as described above. Vaccine dry powder was stored in a desiccator at room temperature until further analysis. Dry powder of AS01_B_/FluzoneQuad was reconstituted in water so that the final expected concentrations of hemagglutinin proteins, DOPC, cholesterol, MPL, QS-21 and sucrose were 60 µg, 1000 µg, 250 µg, 50 µg, 50 µg and 20 mg per 0.5 mL, respectively. The particle size distribution and zeta potential values of liquid (*i*.*e*., before TFFD) and reconstituted AS01_B_/FluzoneQuad vaccine (*i*.*e*., after TFFD) were determined by DLS. The integrity of hemagglutinin proteins was evaluated by SDS-PAGE analysis and intrinsic tryptophan fluorescence spectrometry. The integrity of hemagglutin proteins in the AS01_B_/FluzoneQuad powder was further evaluated by a standard hemagglutination (HA) assay using chicken red blood erythrocytes as previously described [48]. Briefly, 50 *µ*L of the reconstituted vaccine powder was two-fold serially diluted using PBS (10 mM, pH 7.2) in U-bottom 96-well plates (Corning Inc., Corning NY). The samples were then incubated with 50 *µ*L of 1% chicken erythrocyte suspension (Rockland Immunochemicals, Inc., Limerick, PA) in PBS for 30 min at room temperature. HA titers were reported as the reciprocal of the last dilution where HA was observed (*i*.*e*., absence of chicken erythrocyte precipitation) and were expressed in HA units (HAUs)/50 *µ*L.

### 2.5 Preparation and characterizatoin of dry powders of Shingrix vaccine

GSK’s Shingrix was obtained from UT Austin’s Forty Acres Pharmacy. The Shingrix vaccine was prepared by reconstituting the lyophilized powder of VZV-gE antigen with the accompanying suspension of AS01_B_ adjuvant following the Prescribing Information [49]. The vaccine was then frozen to thin-films by single-vial TFF followed by sublimation as described above. It is worth noting that the reconstituted Shingrix vaccine comprises sucrose at sugar to total lipids ratio of 15:1 *w/w*. Single-vial Shingrix powder prepared using TFFD was reconstituted in water before DLS and transmission electron microscopy studies. A Tecnai Spirit Biotwin Transmission Electron Microscope (TEM, ThermoFisher) was used to examine the morphology of the vaccine. Dry powder of Shingrix vaccine was also prepared using shelf freeze-drying. Briefly, one dose (0.5 mL) of reconstituted Shingrix vaccine was gradually cooled from room temperature (~ 21 ± 2°C) to -40°C at a cooling rate of about 2°C/min. The frozen liquid was maintained at -40°C for 1 h in the lyophilizer prior to lyophilization using the same lyophilization cycle for the thin-film frozen samples.

### Statistical analysis

Student’s t-test or one-way analysis of variance (ANOVA) followed by Tukey’s or Dunnet multiple comparison test were performed using GraphPad Prism version 8.0.0 for Windows (GraphPad Software, San Diego, CA). Differences were deemed significant if *p* ≤ *0*.*05*.

## 3. Results and discussion

### 3.1 Dry powders of AS01_B_ adjuvant and AS01_B_/OVA model vaccine

The liposomal formulation is necessary to enhance the safety profile of AS01_B_ immunostimulants. The toxicity of MPL reside in the hydrophobic acyl groups. In AS01_B_, these groups are embedded in the liposome lipid bilayers. Thus, no MPL-induced clinical pyrogenicity is observed after intramuscular injection in human subjects [50]. QS-21-induced damage to the surrounding tissues in the injection site is also circumvented by the irreversible complexation of QS-21 to the cholesterol component of the liposomes [51]. However, the stability of vaccines containing liposomal formulation-based adjuvants is challenged by 248 poor stability of liposomes in liquid form as well as the sensitivity of liposomes to freezing. Herein, we hypothesized that a freeze-insensitive, single-vial presentation dry powder form of AS01_B_-adjuvanted vaccines can be successfully formulated using TFF process followed by lyophilization. Dry powder form is generally more stable than its liquid counterpart [52]. Additionally, dry powders prepared using TFFD have the potential to be delivered *via* noninvasive routes such as intranasal or oral inhalation [35, 40]. Ultimately, single-vial presentation will increase the speed and reduce the cost of vaccine production and vaccination.

Firstly, we explored the effect of TFFD on the physical properties (*i*.*e*., mean particle size, zeta potential and PDI) of AS01_B_ adjuvant alone using sucrose as a stabilizer at the same concentration level present in the final reconstituted Shingrix vaccine (*i*.*e*., 4% *w/v* or lipid : sucrose ratio of 1:16 *w/w*). Ideally, the drying technology should not affect the physical properties, nor the structure of liposomes in order to maintain their adjuvant activity. For instance, the liposome particle size is a key attribute that affects the body’s immune response to the loaded antigens [31]. As shown in **Fig. 1A-B**, the particle size distribution of the AS01_B_ liposomes in the reconstituted AS01_B_ powder was unimodal and identical to that in the liquid AS01_s_. However, the zeta potential values of the AS01_B_ adjuvant decreased after it was subjected to TFFD and reconstitution (*p*<*0*.*01*, **Fig. 1C**), probably due to the strong interaction of sucrose with the liposomes during TFFD [53].

**Fig. 1.**
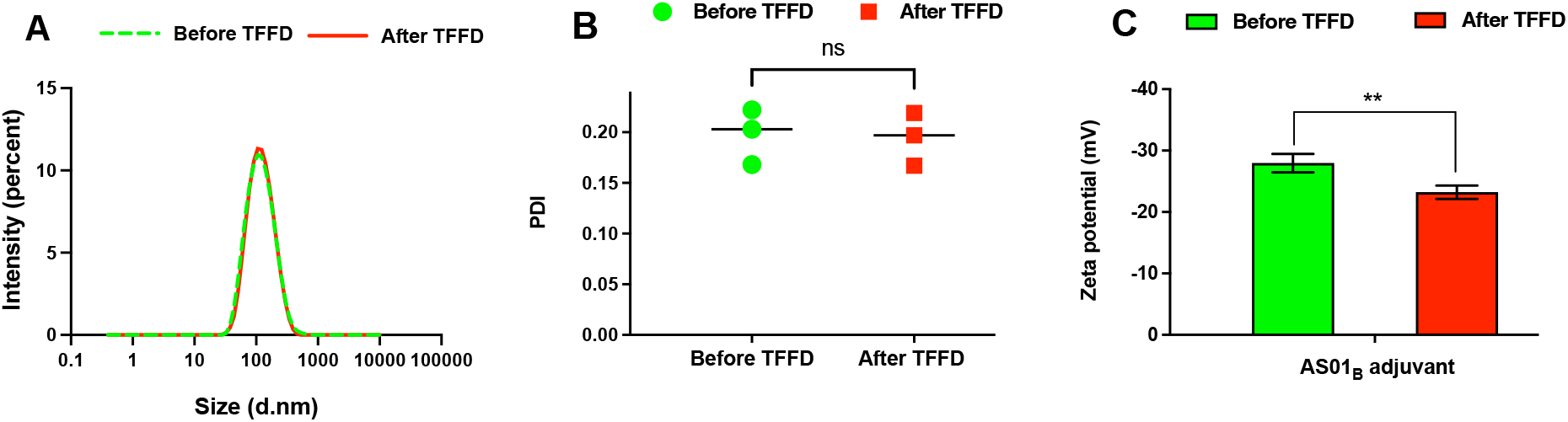
Effect of TFFD and subsequent reconstitution on (A) the particle size distribution, (B) PDI, and (C) zeta potential of AS01_B_ adjuvant. Shown are the properties of freshly prepared liquid AS01_B_ (*i*.*e*., before TFFD) and AS01_B_ subjected to TFFD and reconstitution (*i*.*e*., after TFFD). *ns*: non-significant, \*\**p*<0.01.

Secondly, the applicability of using TFFD to formulate dry powders of AS01_B_-adjuvanted vaccines was investigated using OVA as a model antigen and different stabilizers (*i*.*e*., sucrose, trehalose, and mannitol) at different lipid to stabilizer ratios (*i*.*e*., 1:4, 1:8, 1:15 and 1:30 *w/w*, respectively). Maintaining liposomal integrity during freezing and subsequent drying is challenging [54]. Formation of ice crystals in the internal aqueous compartment or the external aqueous phase during the freezing step can rapture the liposomal lipid bilayers [55]. Saccharides (*e*.*g*., sucrose, trehalose, and mannitol) with many hydroxyl groups are commonly utilized to prevent the aggregation and conformational changes of liposomal lipid bilayers during drying *via* the formation of glassy matrices [56]. High sucrose to lipid ratios (*i*.*e*., 15:1 and 30:1 *w/w*, respectively) have successfully maintained the unimodal particle size distribution of liposomes in the vaccines after drying and reconstitution, unlike low ratios (*i*.*e*., 4:1 and 8:1, **Fig. S1**). Trehalose was successful only when it was incorporated at a ratio of 15:1 *w/w* (trehalose : lipids) while mannitol was unsuccessful at all ratios tested (**Fig. S2 and S3**). Mannitol easily crystallizes during freezing and thus can damage the integrity of liposomal bilayers [13, 57, 58]. Trehalose dihydrate crystallizes upon annealing; however, the dihydrate undergoes dehydration to amorphous anhydrate during drying [59]. Crystallization of lyoprotectants also has deleterious effects on the stability of protein antigens [60]. Sucrose, a non-crystallizing sugar [60], exhibited the optimal protection of the liposomal bilayers against freezing and drying-induced stresses. As depicted in **Fig. S4**, sucrose as a stabilizer in the dry powders of AS01_B_ and AS01_B_-adjuvanted OVA vaccine was amorphous. The stabilizing agent must retain its amorphous solid state during freezing and subsequent drying in order to be effective [59]. The sharp diffraction peaks around 2θ of 27.5°, 31.5°, 45.5° and 56.5° are attributed to the diffraction of NaCl crystals present in the PBS [61]. Lower concentrations of sucrose may have not provided sufficient protection of liposomes against aggregation [55]. Therefore, the AS01_B_/OVA model vaccine containing sucrose at a sugar to lipid weight ratio of 15:1 (*i*.*e*., final sucrose concentration of 4%, *w/v*) was selected for further evaluation and referred to as AS01_B_/OVA. Vaccine dry powders prepared by bulk TFF at drum temperature of -50ºC or -180ºC and subsequently lyophilized also showed unimodal particle size distribution identical to freshly prepared liquid vaccine (*i*.*e*., before TFFD) (**Fig. 5S**). Therefore, the liquid AS01_B_-adjuvanted vaccine can be frozen into thin-films on a small or large scale at a wide range of temperatures.

#### 3.1.1 In-vitro characterization of AS01_B_/OVA vaccine dry powder

As depicted in **Fig. 2A**, the mean particle size of the AS01_B_/OVA model vaccine in sucrose as a stabilizer was maintained after TFFD and reconstitution. However, the zeta potential values of the model vaccine decreased (*p*<0.05, **Fig. 2B**). The integrity of OVA antigen after TFFD was evaluated using SDS-PAGE and intrinsic tryptophan fluorescence spectrometry. SDS-PAGE analysis showed no apparent aggregation or fragmentation of the OVA protein after the model vaccine was subjected to TFFD and reconstitution (**Fig. 2C**). Subjecting the vaccine to TFFD did not affect the wavelength of maximum tryptophan fluorescence emission (*i*.*e*., ʎ_EM_ = 335 nm); however, the intrinsic fluorescence intensity of the three tryptophan residues within the OVA molecule was slightly decreased (**Fig. 2D**). ʎ_EM_ as well as tryptophan fluorescence intensity are extremely affected by the polarity of tryptophan microenvironment and its covalent and noncovalent interactions [62]. Previously, we reported that exposing a tetanus toxoid vaccine adjuvanted with an aluminum salt to TFFD resulted in both a reduction of relative intrinsic tryptophan fluorescence intensity and a slight red shifting of ʎ_EM_ [45]. However, in the current study no shift of ʎ_EM_ was observed, which indicates that the OVA molecules retained some degree of compactness [63]. It is noteworthy that the observed decrease in the relative intensity of intrinsic tryptophan fluorescence could be, in part, attributed to the quenching effect of AS01_B_ (**Fig. S6**).

**Fig. 2.**
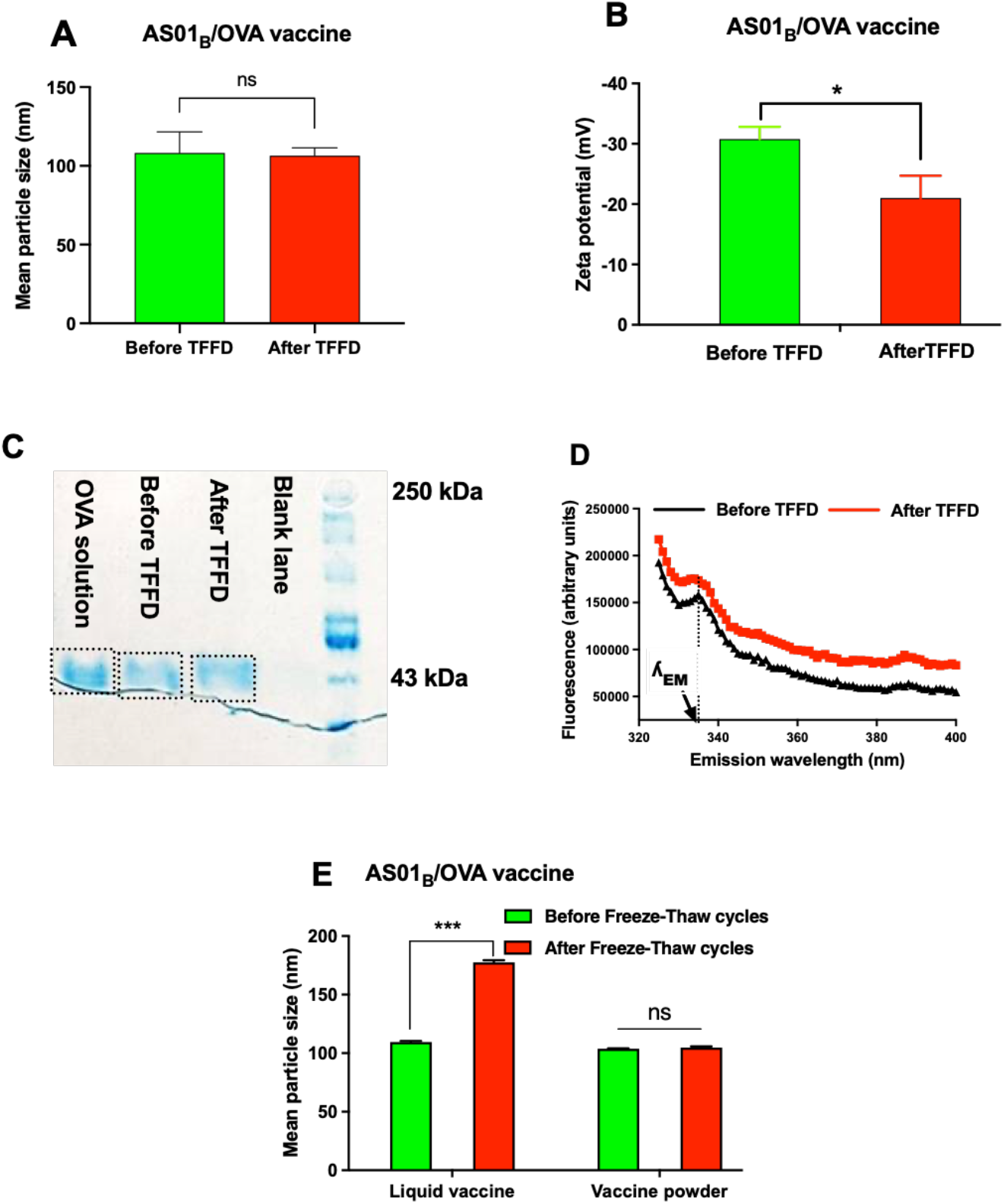
*In-vitro* characterization of AS01_B_/OVA vaccine in liquid (*i*.*e*., before TFFD) and reconstituted from thin-film freeze-dried powder (*i*.*e*., after TFFD). (A) Mean particle size and (B) zeta potential values. (C) SDS-PAGE analysis. (D) Intrinsic tryptophan fluorescence spectra. The final concentration of OVA in both formulations was 100 µg/mL. (E) Mean particle size after three consecutive cycles of freeze-thaw. *ns*: non-significant, \**P*<0.05, \*\**p*<0.01.

Importantly, the mean particle size of AS01_B_/OVA vaccine did not significantly change after the dry powder was subjected to repeated freeze-and-thaw (**Fig. 2E**). On the other hand, subjecting the same liquid vaccine to repeated freeze-and-thaw caused significant aggregation (*p*<0.001). Generally, slow freezing is considered a substantial threat to vaccine potency [64]. It has been estimated that 75-100% of freeze-sensitive vaccines are damaged during transport and storage [16, 64]. Another study reported that 33% and 38% of vaccines have been exposed to freezing temperatures during storage and shipments, respectively, in high-income countries while in low-income countries these ratios are 37.1% and 19.3%, respectively [65]. In addition to cold chain failure, ambient freezing temperatures have been implicated in damage of vaccine potency [66]. Unintentional exposure of vaccines to freezing temperatures not only has resulted in vaccine wastage but also in disease outbreaks [67, 68]. Thus, the development of thermostable or at least freeze-insensitive vaccines is urgently needed. Of course, preparing an AS01_B_-adjuvanted vaccine in dry powder form in a single vial, as opposite to the two-vial presentation of Shingrix vaccine, is also expected to make it cost-effective to transport and store the vaccine.

#### 3.1.2 In-vivo characterization of thin-film freeze-dried AS01_B_/OVA vaccine powder

The ability of a vaccine to induce antigen-specific immune responses is critical for its efficacy [69]. Therefore, an immunization study with the AS01_B_/OVA vaccine, either freshly prepared or reconstituted from thin-film freeze-dried powder, was performed in mice in order to compare their ability in generating OVA-specific immune responses. Female C57BL/6 mice were immunized on day 0 with 5 µg OVA and 135 µg AS01_B_ followed by one booster immunization on day 21. OVA solution in sterile PBS and PBS alone served as controls. Sera and spleens were collected on day 35 upon euthanasia of the mice. As depicted in **Fig. 3A-B**, mice immunized with the AS01_B_/OVA vaccine reconstituted from the thin-film freeze-dried powder showed anti-OVA IgG and IgG1 titers that were not different from that in mice immunized with the same AS01_B_/OVA liquid vaccine that was not subjected to TFFD (p > 0.05). Surprisingly, anti-OVA IgG2a antibodies were not detectable in any of the mice in the study (data not shown). In addition, the splenocytes from mice immunized with the AS01_B_/OVA vaccine reconstituted from the thin-film freeze-dried powder proliferated to a significantly larger extent upon *in-vitro* re-stimulation with OVA than the splenocytes from mice immunized with the same AS01_B_/OVA liquid vaccine that was not subjected to TFFD (**Fig. 3C**). AS01_B_ adjuvant is known to help induce type 1 T helper (Th1) cell immune response [70]. Thus, the OVA-stimulated splenocytes produced predominantly IFN-γ [71] (**Fig. 3D**), with little IL-4 (data not shown). Overall, data in **Fig. 2-3** showed that subjecting the AS01_B_/OVA vaccine to TFFD and subsequent reconstitution did not negatively affect the AS01_B_ liposomes, nor the immunogenicity of the vaccine. Thus, TFFD has made single-vial presentation of AS01_B_-adjuvanted vaccines possible, and the vaccine dry powder is no longer sensitive to accidental freezing.

**Fig. 3.**
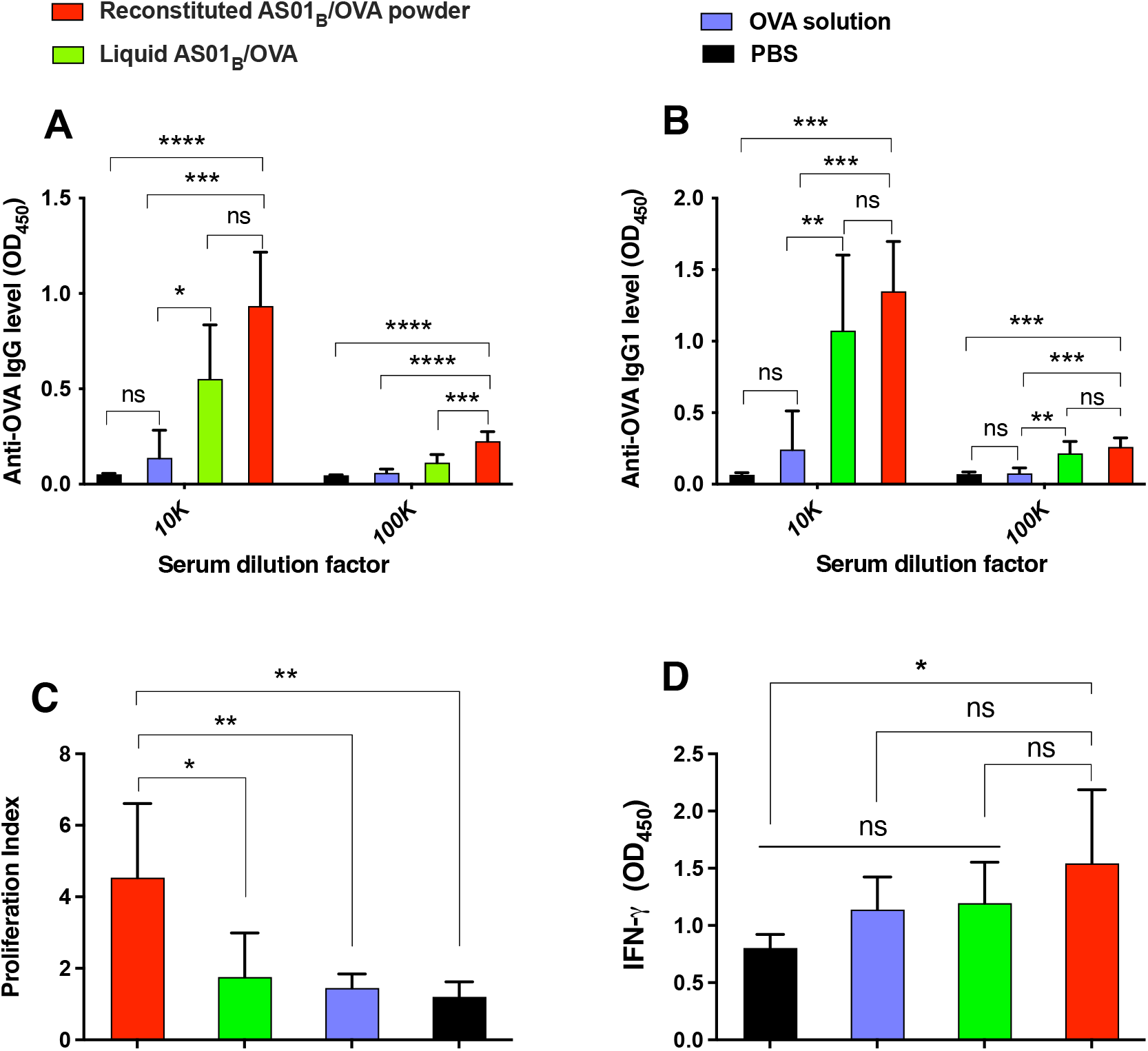
OVA-specific immune responses induced by AS01_B_/OVA vaccine subjected to TFFD. (A) Anti-OVA total IgG and (B) anti-OVA IgG1 antibody levels in mouse sera. (C) Proliferation of splenocytes from immunized mice after *in-vitro* stimulation with OVA. (D) *In-vitro* release of IFN-γ cytokine by mouse splenocytes after *in-vitro* stimulation with OVA. Splenocytes were collected two weeks after the last immunization and cultured in the presence or absence of OVA (10 µg/mL) for 48 h. Data shown are mean 373 ± S.D. (n = 5). *\*p*<*0*.*05, **p*<*0*.*01,***p*<*0*.*001, ****p*<*0*.*0001. ns:* non-significant.

Finally, in a preliminary study, the aerosol performance of the thin-film freeze-dried AS01_B_/OVA vaccine powder containing sucrose and leucine at weight ratios of 71.8% *w/w* and 7.18% *w/w*, respectively, was evaluated using a Next Generation Impactor and a Plastiape RS00 dry powder inhaler at a flow rate of 60 L/min. The mass median aerodynamic diameter of (MMAD) of the powder was ~5.7 µm, suggesting that AS01_B_/OVA vaccine powder may be administered non-invasively to the upper respiratory tract for mucosal immunization [40, 72, 73].

### 3.2 Thin-film freeze-drying of AS01_B_-containing Fluzone Quadrivalent vaccine

Fluzone Quadrivalent is an unadjuvanted, inactivated influenza vaccine that contains the hemagglutinin proteins (60 µg/0.5 mL) of four influenza strains. The vaccine should be stored between 2ºC and 8ºC and must not be frozen [74]. The need for large amount of antigen (*i*.*e*., 15 µg hemagglutinin protein of each viral strain in Fluzone Quadrivalent, and 60 μg of hemagglutinin for each strain in Fluzone High-dose Quadrivalent) to prepare one dose is among the factors that significantly contribute to the limited production of influenza vaccines and can seriously affect the rate of vaccination in the case of a pandemic [75]. One approach to overcome this shortcoming is using an adjuvant [76]. For instance, Audenz, an FDA-approved influenza A (H5N1) vaccine adjuvanted with MF59, contains only 7.5 µg hemagglutinin/0.5 mL. Adjuvants can also enhance the immunogenicity of influenza vaccines in elderly who are less responsive to vaccines [77]. Although the world is concerned about the threat of influenza pandemics [78], only two adjuvants (*i*.*e*., MF59 and AS03) are used in FDA-approved influenza vaccines [1]. AS01_B_ may be potentially used for future development of seasonal or universal pandemic flu vaccines. Therefore, in this study, we investigated the feasibility of preparing an AS01_B_-adjuvanted influenza vaccine dry powder by TFFD. For this purpose, an AS01_B_-adjuvanted FluzoneQuad (AS01_B_/FluzoneQuad) vaccine was prepared by mixing 0.5 mL of in-house prepared AS01_B_ adjuvant with an equal volume of FluzoneQuad. Sucrose was employed as a stabilizer at sugar to total lipid ratio of 15:1, *w/w*, and the liquid vaccine was then subjected to TFFD.

As depicted in **Fig. 4A-B**, the mean particle size of the AS01_B_/FluzoneQuad vaccine was generally preserved after it was subjected to TFFD, but the zeta potential was decreased. SDS-PAGE analysis showed no apparent aggregation or fragmentation of hemagglutinin proteins (**Fig. 4C**). This observation was further confirmed by investigating the intrinsic tryptophan fluorescence of the AS01_B_/FluzoneQuad vaccine after subjected to TFFD and reconstitution. The fluorescence spectra of the liquid AS01_B_/FluzoneQuad vaccine (*i*.*e*., before TFFD) and the vaccine reconstituted from the thin-film freeze-dried powder (*i*.*e*., after TFFD) were similar (*i*.*e*., no shift in *λ*_*EM*_ or decrease in fluorescence intensity) (**Fig. 4D**). Importantly, the hemagglutination activity of the AS01_B_/FluzoneQuad vaccine in terms of HA titer was preserved after it was subjected to TFFD and subsequent reconstitution (**Fig. 4E**). For unknown reasons, the addition of AS01_B_ into the FluzoneQuad vaccine led to an increase in the HA titer of the vaccine (**Fig. 4E**). Finally, unlike the AS01_B_/FluzoneQuad liquid vaccine (**Fig. 4F**), the particle size distribution of the AS01_B_/FluzoneQuad in the thin-film freeze-dried powder remained unchanged after the powder was subjected to three cycles of repeated freeze-and-thaw followed by reconstitution (**Fig. 4F-G**). Taken together, data in **Fig. 4** demonstrate that it is feasible to prepare a dry powder form of an influenza vaccine adjuvanted with AS01_B_ by using TFFD. The new vaccine composition and TFFD technology represent a step forward in preparing the world for future influenza pandemics.

**Fig. 4.**
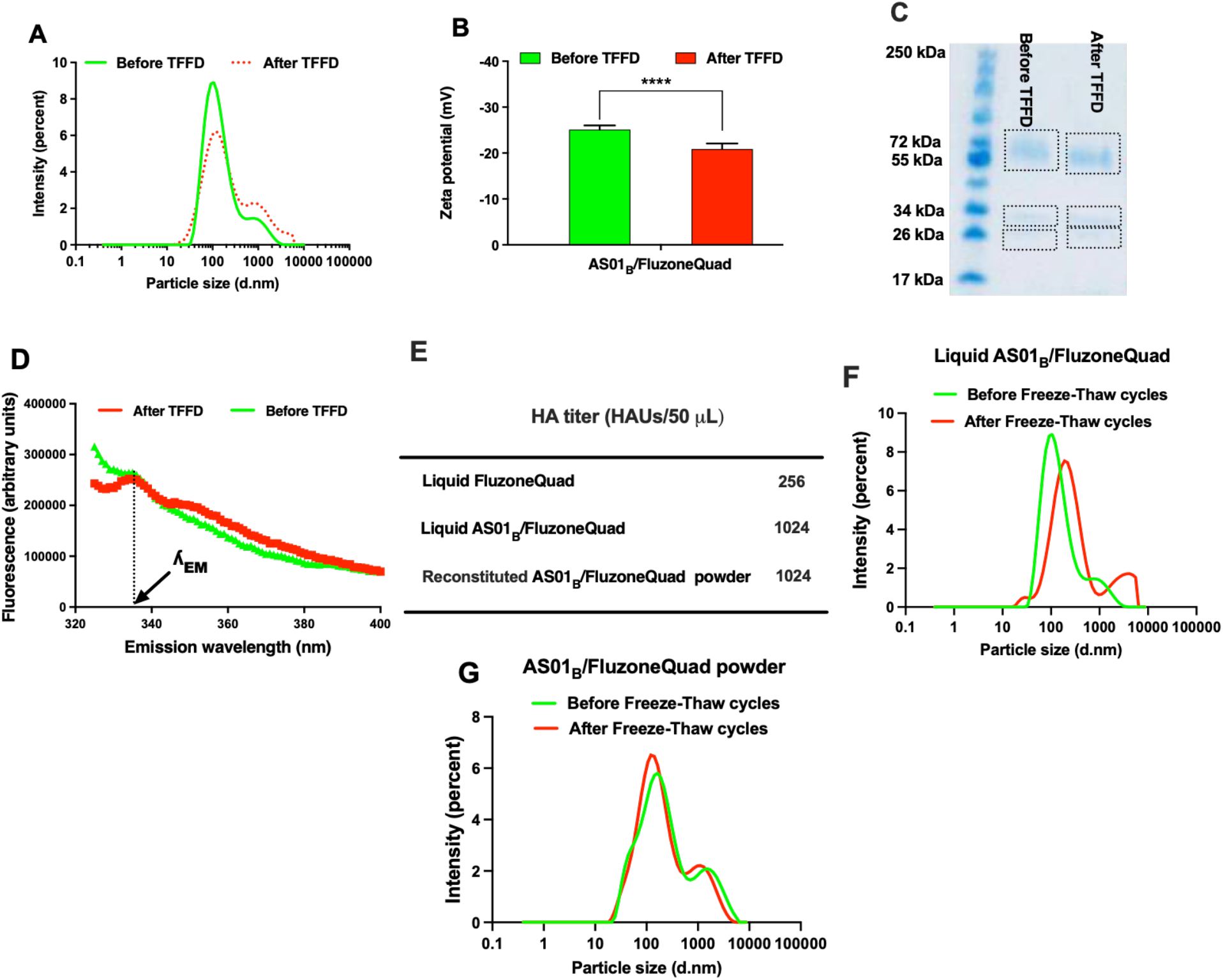
Thin-film freeze-drying of AS01_B_-adjuvanted Fluzone Quadrivalent vaccine. (A) Particle size distribution and (B) zeta potential values of liquid (*i*.*e*., before TFFD) and reconstituted (*i*.*e*., after TFFD) vaccine. (C) SDS-PAGE image showing the hemagglutinin proteins in liquid vaccine as well as the vaccine reconstituted from the dry powder. (D) Intrinsic tryptophan fluorescence spectra. (E) Hemagglutination titers of the original Fluzone Quadrivalent vaccine (*i*.*e*., liquid FluzoneQuad), liquid AS01_B_/FluzoneQuad and AS01_B_/FluzoneQuad reconstituted from the dry powder prepared using TFFD (*i*.*e*., reconstituted AS01_B_/FluzoneQuad). The hemagglutination assay was repeated twice with the same results. (F-G) Particle size distribution of AS01_B_/FluzoneQuad vaccine after the liquid vaccine (F) or the dry powder (G) was subjected to three cycles of freeze-thaw. *ns*: non-significant (*p>0*.*05*), \*\*\**p*<*0*.*01*.

### 3.3 Thin-film freeze-drying of Shingrix vaccine

Shingrix is in a two-vial presentation; the AS01_B_ adjuvant is provided as a liquid suspension in a vial, which is used to reconstitute the VZV-gE antigen as a lyophilized powder in a separate vial immediately before administration. We tested the feasibility of preparing a single vial dry powder form of the Shingrix vaccine (*i*.*e*., the VZV-gE antigen and AS01_B_ adjuvant in a single vial in dry powder) by TFFD. Because the VZV-gE lyophilized powder already contained sucrose, no additional stabilizer was added before the vaccine was subjected to TFFD. As a control, the same vaccine was also subjected to conventional shelf freeze-drying. As depicted in **Table 1** and **Fig. 5A**, the particle size distribution, mean particle size, and zeta potential of Shingrix vaccine reconstituted from the thin-film freeze-dried powder were not different from the original liquid Shingrix vaccine. These results were confirmed by the transmission electron microscopy images of the vaccine reconstituted from the thin-film freeze-dried powder (**Fig. 5B**). The images showed uniform particles without apparent aggregation (**Fig. 5B**). In contrast, subjecting the same Shingrix vaccine to shelf freeze-drying led to a significant increase in the particle size upon reconstitution (**Table 1** and **Fig. 5A**), indicating why the Shingrix vaccine is currently marketed in a two-vial presentation. The TFFD technology can potentially enable the packaging of the two components of the Shingrix vaccine in a single vial as a dry powder. Single-vial presentation can minimize the cost and complexity of manufacturing and increase the rate of vaccination since it is more convenient for health care providers [79]. The developed vaccine powder will likely be more stable than its liquid counterpart and freeze-insensitive and thus can potentially reduce vaccine wastage [52].

**Fig. 5.**
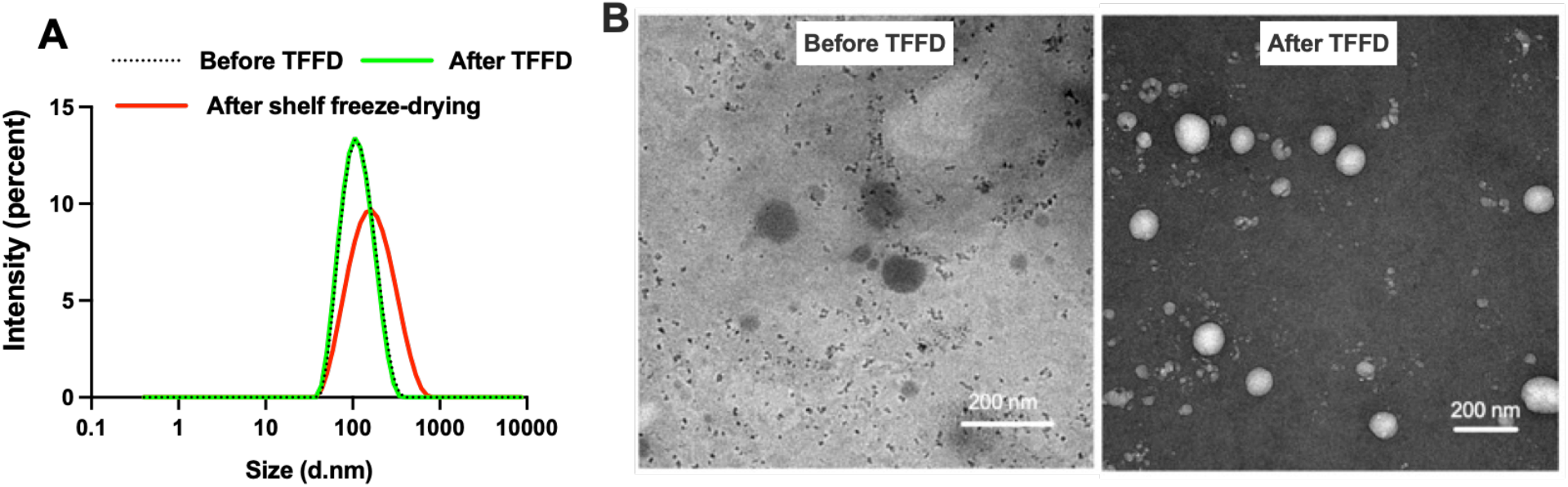
Characterization of Shingrix dry powders prepared using TFFD or shelf freeze-drying. (A) Particle size distribution of liquid Shingrix vaccine (*i*.*e*., before TFFD) and Shingrix vaccine reconstituted from dry powders prepared using TFFD (*i*.*e*., after TFFD) or shelf freeze-drying. (B) Representative TEM images of liquid Shingrix vaccine and Shingrix vaccine reconstituted from dry powder prepared using TFFD.

**Table 1.** Mean particle size, PDI, and zeta potential values of liquid Shingrix vaccine (*i*.*e*., before TFFD) and Shingrix vaccine reconstituted from dry powders prepared using TFFD (*i*.*e*., after TFFD) or shelf freeze-drying. Data are mean ± S.D. (n = 3). *ns:* non-significant, *\*\*p*<*0*.*01* compared to the original liquid Shingrix vaccine.

| Vaccine | Mean particle size<br>(nm) | PDI | Zeta potential<br>(mV) |
| --- | --- | --- | --- |
| Liquid Shingrix vaccine | 105 ± 1 | 0.12 ± 0.01 | -14 ± 2 |
| Shingrix vaccine after TFFD | 105 ± 0.3<sup>ns</sup> | 0.19 ± 0.02<sup>**</sup> | -12 ± 1 |
| Shingrix vaccine after shelf freeze-drying | 141 ± 2<sup>**</sup> | 0.23 ± 0.01<sup>**</sup> | NA |

## Conclusion

Thin-film freeze-drying enabled the preparation of dry powders of the AS01_B_ liposomal adjuvant and AS01_B_-adjuvanted vaccines using a low concentration of a single stabilizing agent. The liposome integrity and vaccine immunogenicity were maintained in the dry powders. The resulted AS01_B_-adjuvanted vaccine dry powder is in a single-vial presentation, is not sensitive to freezing, and may be stored cold chain-free.

## Acknowledgments

This work was supported by Sponsored Research Agreements and Technology Validation Agreements from TFF Pharmaceuticals Inc. (to ROW and ZC). KA is supported in part by a fellowship (GM 1105) from the Egyptian Ministry of Higher Education.

## Conflicts of interest

ZC and ROW report financial support by TFF Pharmaceuticals, Inc. ZC reports a relationship with TFF Pharmaceuticals, Inc. that includes: equity or stocks and funding. ROW reports a relationship with TFF Pharmaceuticals, Inc. that includes: consulting or advisory, equity or stocks, and funding. HX report a relationship with TFF Pharmaceuticals, Inc. that includes: consulting or advisory. ZC and ROW have a patent “Dry solid aluminum adjuvant-containing vaccines and related methods thereof” pending to TFF Pharmaceuticals, Inc. ZC, ROW, KA and HX have a patent “Dry liposome adjuvant-containing vaccines and related methods thereof” pending to UT Austin. ZC, ROW and KA have a patent “Dry liposome formulations and related methods thereof” pending to UT Austin.

## Supplementary information

**Fig. S1.**
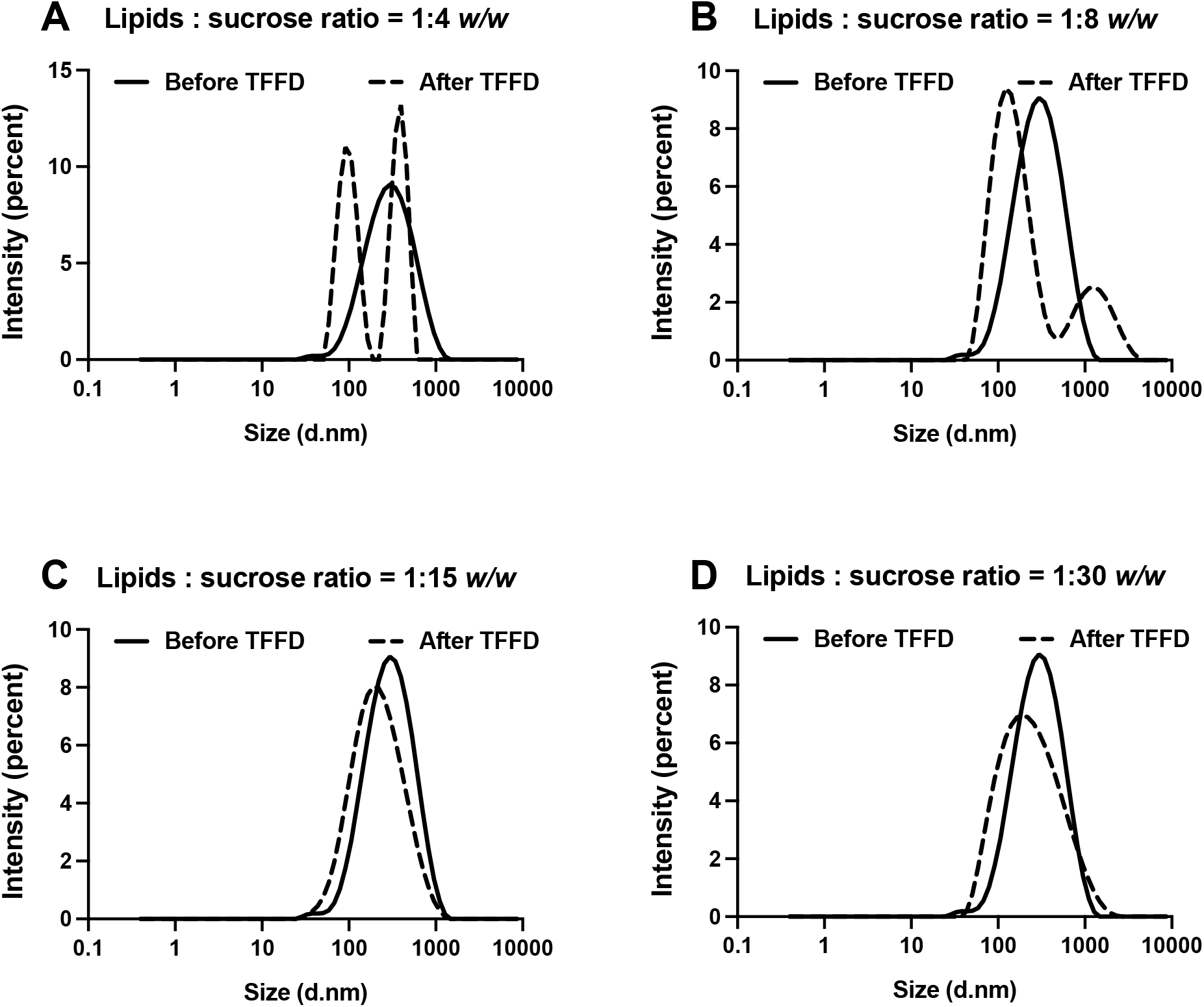
Effect of different lipids to sucrose ratios on the particle size distribution of AS01_B_/OVA model vaccine subjected to TFFD and reconstitution.

**Fig. S2.**
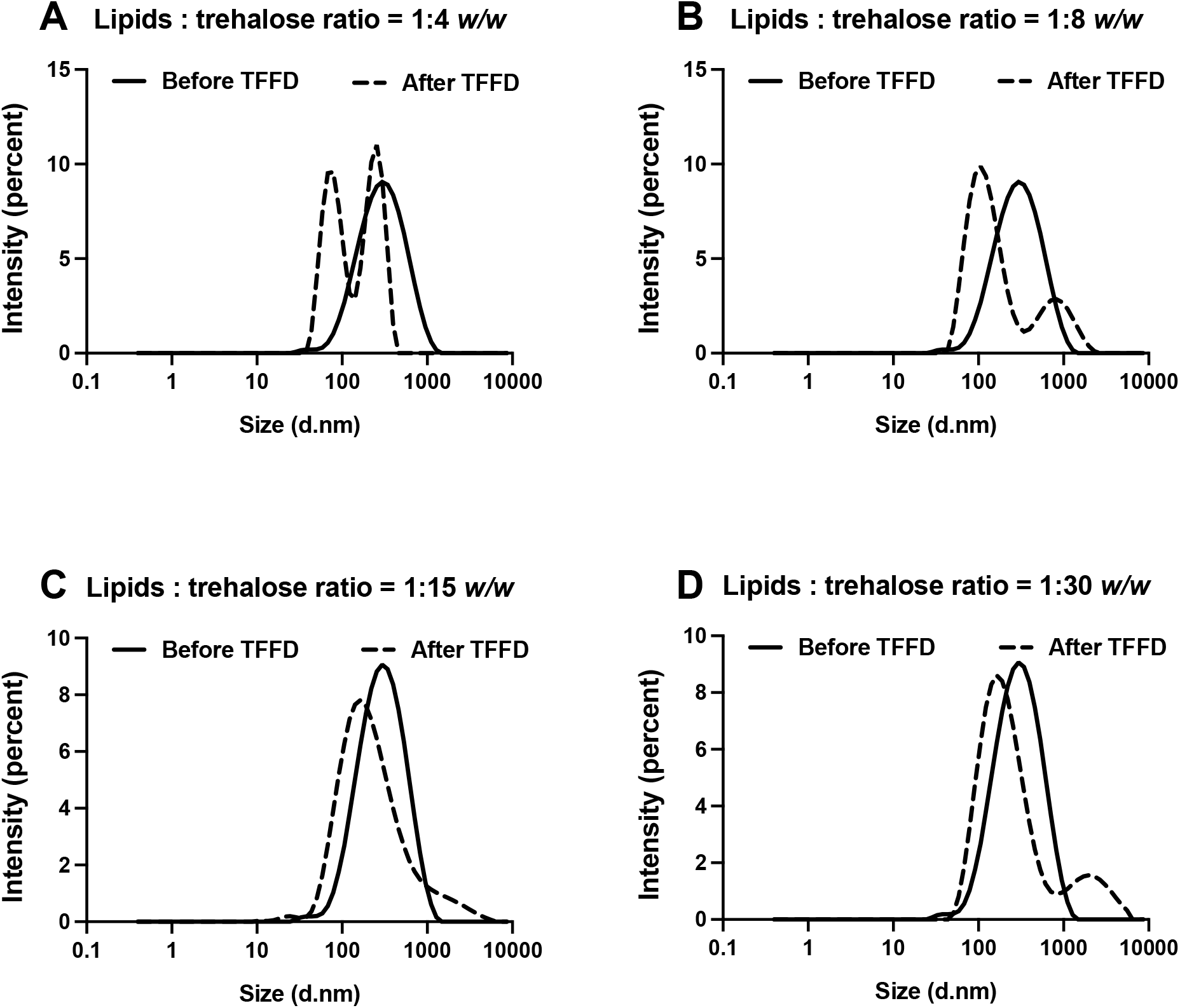
Effect of different lipids to trehalose ratios on the particle size distribution of AS01_B_/OVA model vaccine subjected to TFFD and reconstitution.

**Fig. S3.**
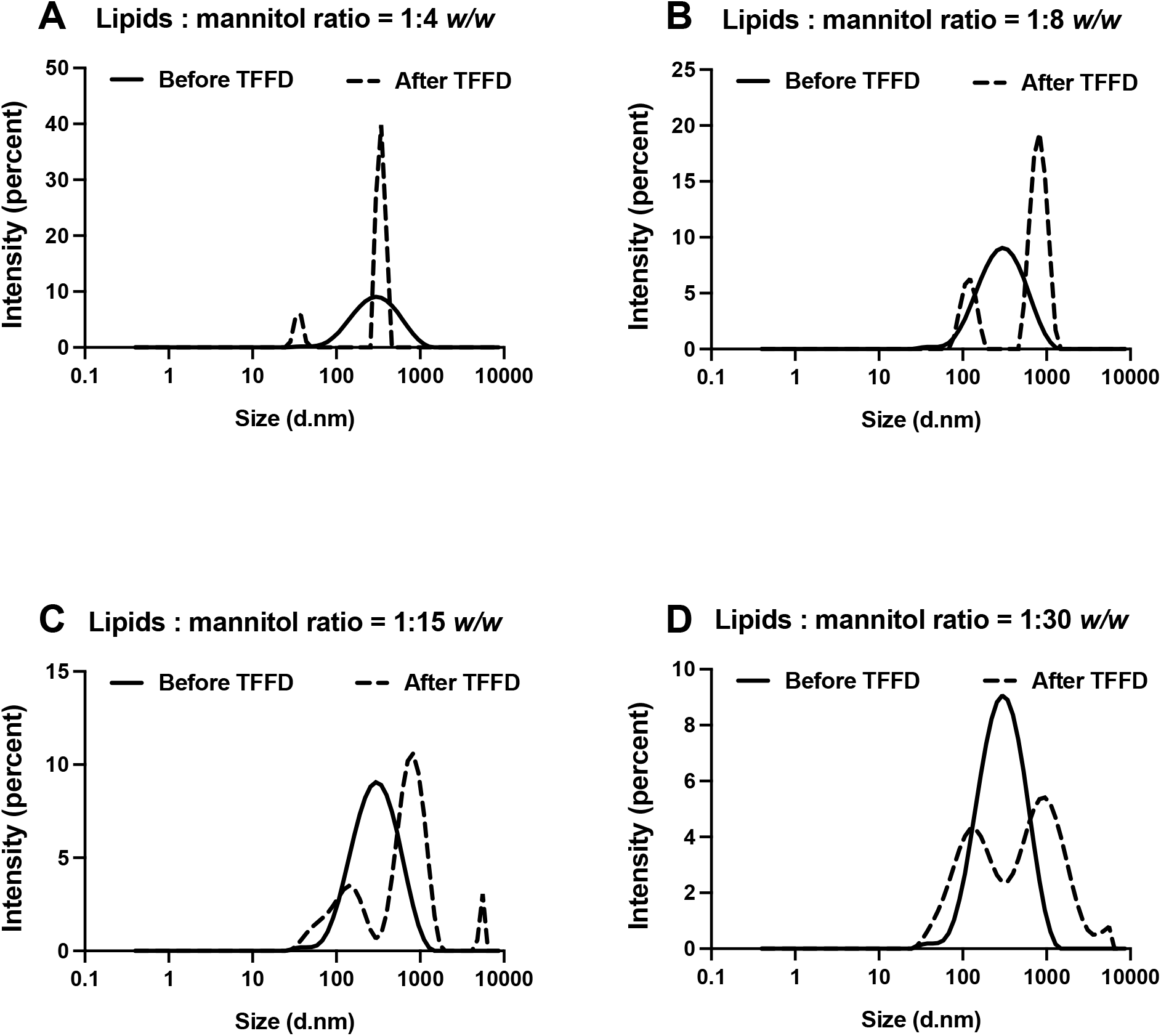
Effect of different lipids to mannitol ratios on the particle size distribution of AS01_B_/OVA model vaccine subjected to TFFD and reconstitution.

**Fig. S4.**
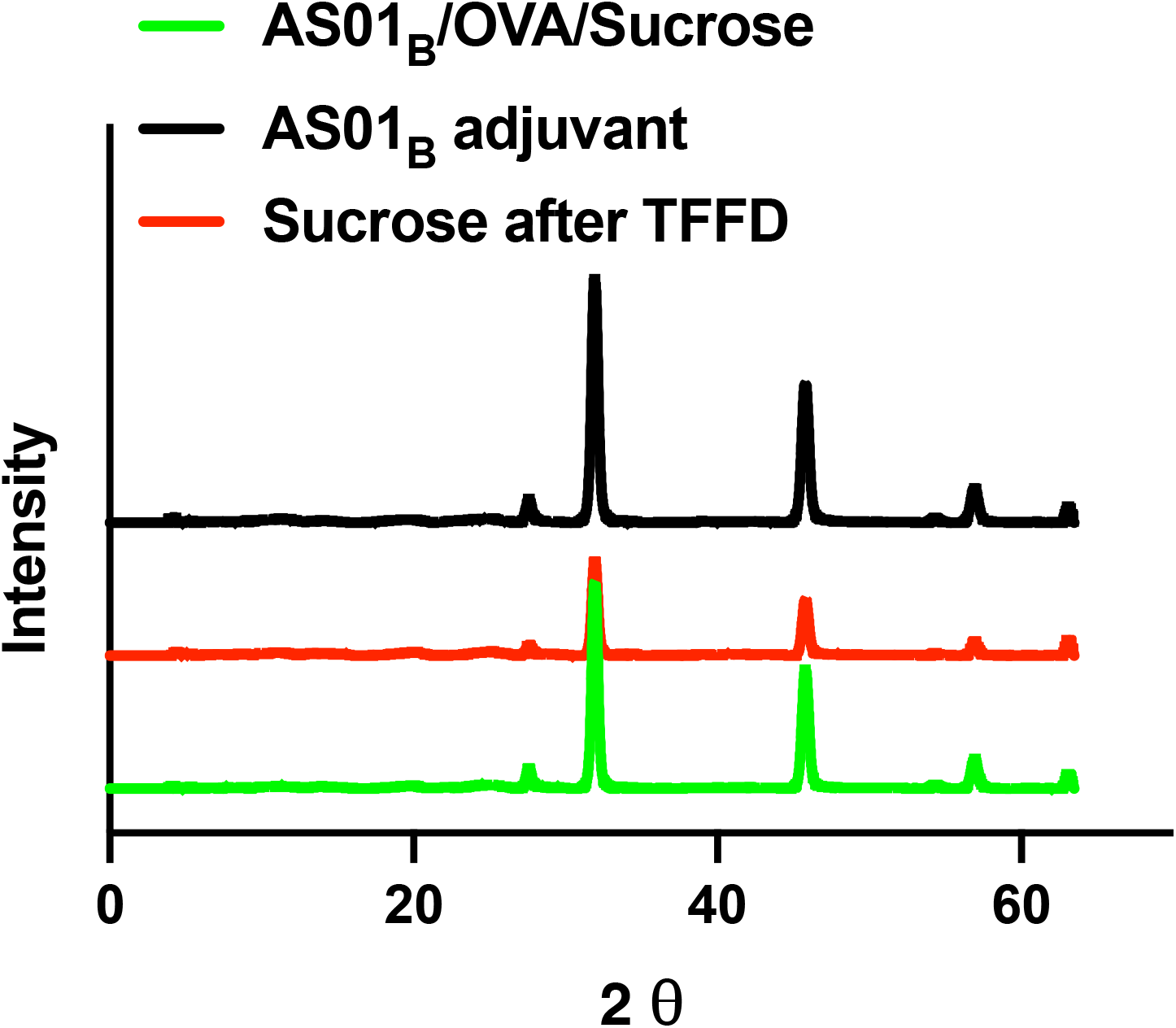
X-ray diffraction patterns of AS01_B_/OVA vaccine and AS01_B_ adjuvant dry powders. The AS01_B_/OVA vaccine contains sucrose as a stabilizer at sucrose to lipid ratio of 15:1 *w/w* (*i*.*e*., 4% *w/v)*. Sucrose was dissolved in PBS at 4% *w/v* then was dried using TFFD.

**Fig. S5.**
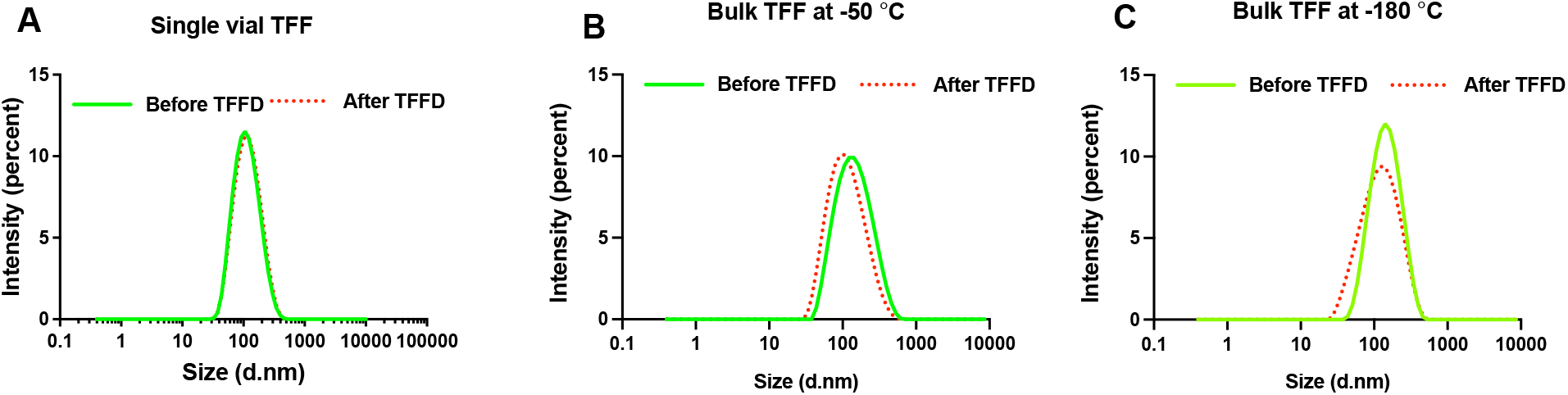
Effect of thin film freezing (TFF) technique (single vial *vs* bulk) and TFF temperature (−50ºC *vs* - 180ºC) on the particle size distribution of AS01_B_/OVA vaccine comprising sucrose at a ratio of 1:15 *w/w* (lipids : sucrose). (A) Liquid AS01_B_/OVA/Sucrose vaccine was dropped onto the inner wall of a cryogenically cooled vial bottom using a syringe (*i*.*e*., single vial TFF). (B) and (C) Liquid AS01_B_/OVA/Sucrose was dropped onto a rotating, cryogenically cooled stainless-steel drum having a temperature of -50°C or -180°C, respectively, to form frozen thin-films.

**Fig. S6.**
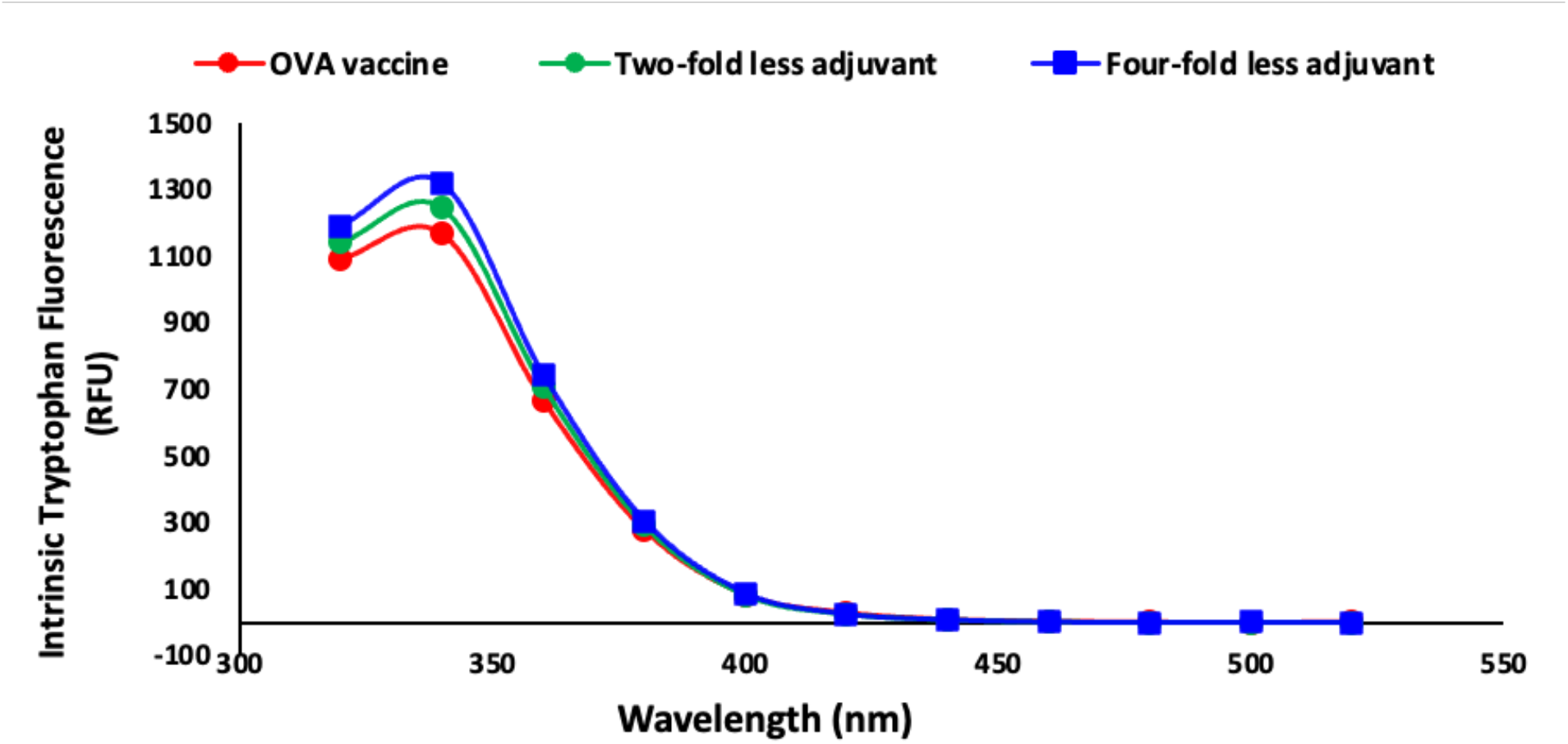
The slight quenching of intrinsic tryptophan fluorescence by the AS01_B_ adjuvant. OVA vaccine containing 1x adjuvant showed the least tryptophan fluorescence intensity followed by the vaccine containing two-fold less AS01_B_ adjuvant, while the vaccine containing four-fold less adjuvant showed the highest tryptophan fluorescence intensity.

